# Creating detailed metadata for an R Shiny analysis of circadian behavior sequence data

**DOI:** 10.1101/2021.07.16.452645

**Authors:** Julien Colomb, York Winter

**Affiliations:** Department of Biology, Humboldt Universität zu Berlin, Berlin, Germany; Exzellenzcluster NeuroCure, Charité, Berlin, Germany

**Keywords:** Home cage scan, mus musculus, rodent, automatic, machine learning, multidimensional analysis

## Abstract

Automated mouse phenotyping through the high-throughput analysis of home cage behavior has brought hope of a more effective and efficient method for testing rodent models of diseases. Advanced video analysis software is able to derive behavioral sequence data sets from multiple-day recordings. However, no dedicated mechanisms exist for sharing or analyzing these types of data. In this article, we present a free, open-source software actionable through a web browser (an R Shiny application), which can perform state-of-the-art multidimensional analysis of home cage behavioral sequence data. The software aligns time-series data to the light/dark cycle, and then uses different time windows to produce up to 162 behavior variables per animal. It prevents p-hacking by providing an analysis that uses a principal component analysis strategy, while also representing the behavior graphically for further explorative analysis. A machine-learning approach was implemented, but it proved ineffective at separating the experimental groups.

The software requires spreadsheets that provide information about the experiment (i.e., metadata), thus promoting a data management strategy that leads to FAIR data production. This encourages the publication of some metadata even when the data are kept private. We tested our software by comparing the behavior of female mice in videos recorded twice at 3 and 7 months in a home cage monitoring system. This study demonstrated that combining data management with data analysis leads to a more efficient and effective research process.

## INTRODUCTION

Any attempt to identify the behavioral phenotype of an animal can be a highly tedious undertaking. Animal behavior depends heavily on many variables, which are sometimes uncontrollable, such as general health, age, animal care, sex, environmental factors (pre- and post-natal), housing conditions, environmental stress (including from the experimenter), and diet [Van Meer and Raber, 2005]. Therefore, the research community has been searching for high-throughput technologies and methods that can not only phenotype numerous animals through computer automation and with low effort from the experimenter, but also be applied without the experimenter interacting with the animal. Behavioral analysis of video-captured home cage behavior could potentially be an effective and efficient method for characterizing rodent models of diseases. Because analyzing the behavior of animals under crowded conditions in group housing remains difficult (see Bains et al. [2018a] for a review), the most widely used approach is to record animals’ behavior in individual cages.

Currently, various technical solutions can provide a detailed analysis of single-housed mouse behavior sequences. These include proprietary systems such as the HomeCageScan (HCS) software (Cleversys, Steele et al. [2007]), and the phenorack system (Viewpoint S.A., France Bains et al. [2018b]), as well as an open source solution [Jhuang et al., 2010]. These software solutions use video analysis to assign one behavior to each video frame (using a short video sequence as the input). The primary data output is a sequence of behavior states. The number of different behaviors recognized varies between the solutions. To simplify the analysis or compare software accuracy, the number of behavior categories can be reduced [Jhuang et al., 2010, Adamah-Biassi et al., 2013, Luby et al., 2012a, Steele et al., 2007]. In addition to the raw behavior sequence data, the HCS software, as one example, may create summaries of the time spent performing each recognized behavior, as well as of the distance traveled (horizontally) for time intervals from minutes to several days.

While much effort has been spent on developing software that automatically tracks behavioral motives over time, very little effort has been invested into the analysis of the data produced. For instance, the amount of data recorded and the time that the experiment starts sometimes vary between animals for technical reasons, but such discrepancies were usually not corrected for. While primary sequences contain up to 45 different behavioral categories recorded over 24 hours, published accounts have mostly reported analyses conducted after data were pooled into only two categories and one time window (approx. 24 h in Steele et al. [2007], 2.5h before feeding time in Luby et al. [2012a]). Consequently, such analyses would detect a difference in overall activity, but not in activity rhythm [Tobler et al., 1996]. On the other hand, a highly detailed analysis of each behavior would lead to a very high number of variables, which would require either a multivariate analysis to avoid harking and false positives or a preliminary experiment to identify the variables of interest *a priori* [Damrau et al., 2021].

Multidimensional approaches have been used previously to separate experimental groups. Steele et al. [2007] used a two-out validation strategy with an L1-regularized logistic regression; specifically, they trained a model on half of the data and then used the model to predict the grouping in the remaining data. This allowed them to discriminate between sick and healthy individuals from the video data well before the appearance of traditionally used symptoms. Another study [Bains et al., 2018a] performed a canonical discriminant analysis to select the behavior variables that best separated groups (animal behaviors were monitored manually).

Currently, no repository exists for home cage monitoring data of animal models of disease. For this study, we obtained only derived data (hourly-binned data exports Steele et al. [2007]), because the raw data had not been saved. This restricted our use of meta- and comparative analyses.

In this article, we present an integrated solution for the analysis and management of home cage video monitoring data. We propose a simple metadata schema in the form of spreadsheets that allow for a flexible structure of the data. The data become computer-readable, a first and critical step toward the production of FAIR (findable, accessible, interoperable, and reusable) open data [Group, 2014]. In addition, we provide a pack of open-source R scripts and R Shiny applications (apps) that can analyze such FAIR data. The software synchronizes the data with light/dark conditions, discards data outside of the time window common to all subjects, and then calculates summary statistics for each behavior for several interesting time windows (defined along the day/night cycle of animal activity). Our software can run a multidimensional analysis that tests whether different experimental groups can be distinguished (using the first component after a principal component analysis [PCA] or based on a machine-learning strategy). It also provides plots of hourly activities for explorative analysis.

In this study, we used unpublished data obtained in Berlin, as well as previously published data obtained from Andrew Steele’s lab, for testing the software and for further development. In particular, we compared the behavior profile of animals monitored twice at 3 and 7 months of age. Because of the differences in age and experience, we expected a change in behavior, which our analysis was in fact able to detect.

## MATERIALS AND METHODS

### Animal testing

The natural behavior of single mice within a home cage, unaffected by an experimenter, was video-recorded from a side view. Animals were singly housed for approximately 23 h in a regular home cage (EU type II) without additional enrichment (to avoid the detection of artifacts on video), but with free access to food and water. The videos were analyzed to classify the single behavior shown on each frame using the HCS software package (CleverSys Inc., USA).

### Software and data availability

We used Rstudio and GitHub to develop the open source software (MIT licensed) as well as to organize its development and version control (www.github.com/jcolomb/HCS_analysis). Github issues were used to archive some discussions held with the CleverSys staff and data providers (Andrew Steele). Different milestones of the development were and will be archived on Zenodo to assure long-term preservation of the software (DOI:10.5281/zenodo.1162721). Data were added to the repository. Different text files available with the software describe and document the use of the two apps, details of the analysis algorithms, and ways to expand the analysis. A readme file explains how to navigate them.

### Main dependencies

The software was built on R resources [R Core Team, 2020]. This work would not have been possible without the tidyverse environment [Wickham, 2019], packages for interactive processing [Pedersen et al., 2020, Chang et al., 2020, Sievert et al., 2020], statistical analysis [Meyer et al., 2019, Breiman et al., 2018, Helwig, 2018, Park and Hastie, 2018, Harrell, 2020] and graphical interfaces [Murrell, 2020, Auguie, 2017, Sievert et al., 2020]. It also depended on the osfr package, which was still in development [Wolen and Hartgerink, 2020] and loaded via the devtools package [Wickham et al., 2020]. We used the env package [Ushey, 2020] to dock the project.

### Metadata structure

The metadata was structured in different files to avoid having to provide the same information multiple times (Figure 1). Each experiment was described in the master metadata file available online (https://osf.io/myxcv/). We expanded the RADAR descriptive metadata schema [Kurze et al., 2017] to create the structure of the master metadata (Table 1). The information entered in that file was made openly available even if the data was not. In addition to the generic entries from RADAR, the file contained information about the path to the other three metadata files - the experiment (one row per test provides details about the animal and the experiment), lab (conditions such as light conditions are given), and identifiers metadata file - and the data folder, as well as information about the software used to acquire and analyze the video.

**Figure 1.**
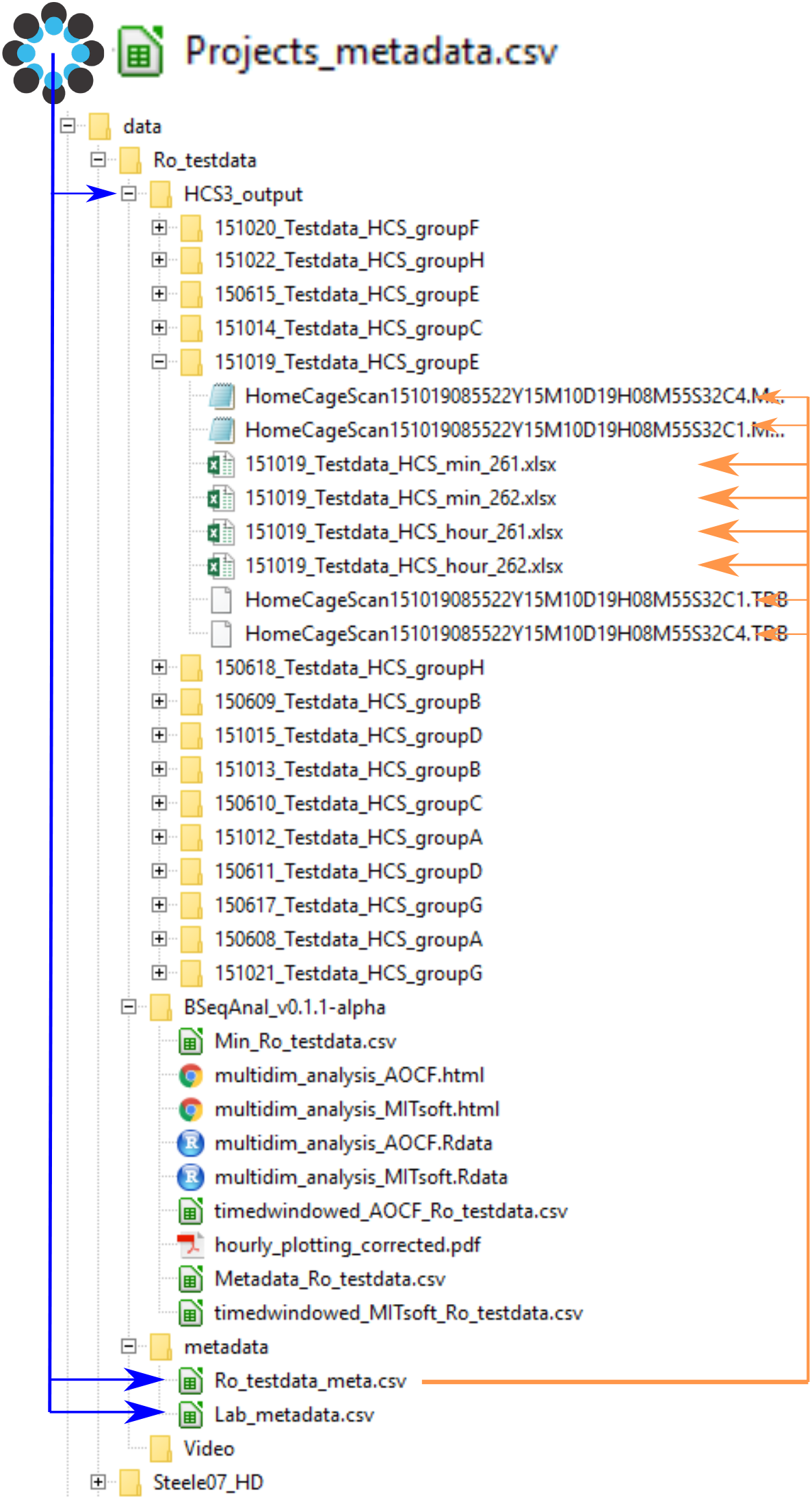
Data and metadata structure. The master project metadata file links the address of the metadata files and the data folder. The experiment metadata file links to each data file (for clarity, only one folder is shown here). The format of the data is either .xlsx summary files (min or hour) or the HCS output files .mbr (behavior sequence) and .tbd (position; note that the software does not read that file). By reading the master file, the computer can determine the path to every data file. Upon analysis, the software creates a new folder indicating the software’s name and version. Its reports are saved there, while derived data files are saved in a folder named after the software name but not its version.

**Table 1.**
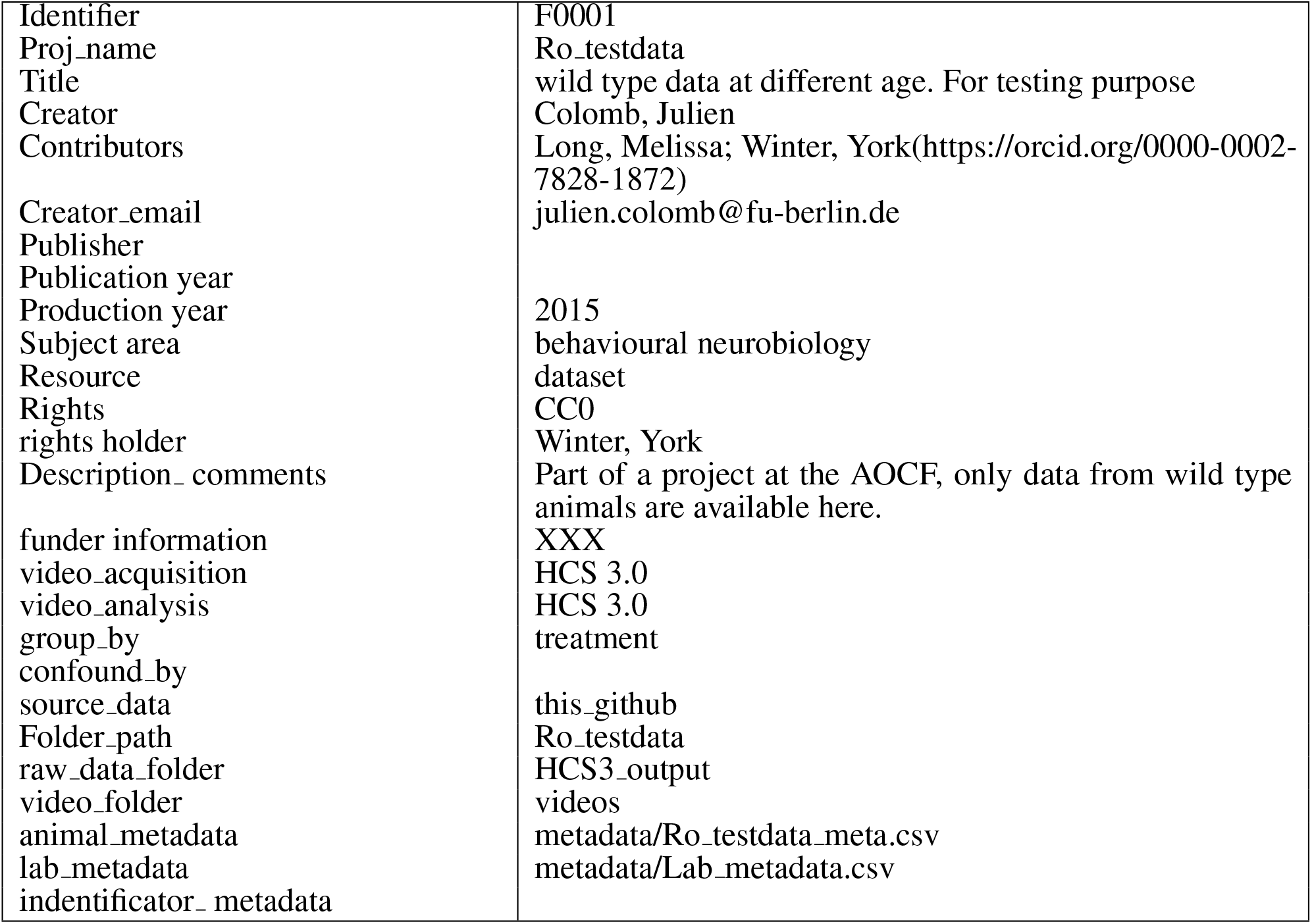
Master metadata information.

### Metadata and data registration

We have provided a detailed manual that describes the relatively complex metadata creation process (see the readme file), and also a Shiny app for testing the quality of the metadata entered by users (available at analysis/Shiny testanduploaddata/) and pushed it to OSF. We followed the manual and push metadata from a different experiment performed in Berlin.

The master metadata file was deposited on the open science framework storage at “https://osf.io/myxcv/” via the Shiny app. We chose this solution not only because we could read and update it directly from R, but also because it was version controlled (i.e., misuse will not have serious consequences). This file indexed all experiments that were analyzed with the software, but the deposition of the actual data remained independent and optional. The analysis software could access data locally (as in the example provided) or on the web via the HTTP protocol. We used the Github repository as one practical example.

### Data analysis

The detailed process of the analysis can be read directly from the commented code and readme file available on Zenodo and GitHub. Variables can be entered in the master noshiny.r file or via the Shiny app, and the master shiny.r file is then processed. The second tab in the Shiny app plots hourly summary data by running the “plot5 hoursummaries.r” code. A brief description of the software procedure is provided below.

#### Overview

The analysis software automatically reads the master metadata file on OSF. When the user specifies the project to be analyzed, the software (1) reads the metadata associated with the project and creates a minute summary file from the indicated primary data file; (2) behavior categories (see Table 2) are pooled and the software creates time windows before calculating a value for each behavior category for each time window. Some data might be excluded from the analysis at this point, following the label indicated in the experiment metadata. The software then performs multidimensional analyses on the latter data to plot them (3) as well as to test whether experimental groups can be distinguished (4). The user can choose which time window to incorporate with step 3. The analysis involves a random forest (RF) analysis for identifying the variables that exhibit the largest differences among the different groups of mice. Next, an independent component analysis (ICA) is performed on these 8–20 variables and the first three components are plotted in an interactive 3D plot. Independently, the next part of the software runs a PCA and examines the first principal component for statistically different results in the groups using a nonparametric test. Then, it may run a machine-learning algorithm on the data. Validation of the latter results is conducted through a non-exhaustive two-out validation technique as in Steele et al. [2007] if the sample size per group is below 15, or otherwise through a test data set.

**Table 2.** The initial 45 categories were pooled into 18 and 10 categories.

| Original_HCS | Berlin_category | Jhuang_category |
| --- | --- | --- |
| Travel.m. | Distance_traveled | Distance_traveled |
| ComeDown | ComeDown | Rear |
| RearUp | Rearup | Rear |
| Turn | Walk | Walk |
| Stretch | Stretch | Rear |
| HangCudl | Hang | Hang |
| HangVert | Hang | Hang |
| CDfromPR | ComeDown | Rear |
| CDtoPR | Rearup | Rear |
| RUfromPR | Rearup | Rear |
| RUtoPR | Rearup | Rear |
| LandVert | Hang | Rear |
| WalkLeft | Walk | Walk |
| WalkRght | Walk | Walk |
| Stationa | Immobile | Rest |
| Drnk.S1. | Drink | Drink |
| Eat.Z1. | Eat | Eat |
| Jump | Jump | Unknown_behavior |
| Unknown | Unknown | Unknown_behavior |
| HVfromRU | Hang | Hang |
| HVfromHC | Hang | Hang |
| ReptJump | Jump | Unknown_behavior |
| Circle | Walk | Walk |
| Dig | Digforage | Unknown_behavior |
| Forage | Digforage | Unknown_behavior |
| Pause | Immobile | Micro_move |
| Urinate | Unknown | Unknown_behavior |
| Groom | Groom | Groom |
| Sleep | Immobile | Rest |
| Twitch | Twitch | Micro_move |
| Arousal |  |  |
| Awaken | Awaken | Micro_move |
| Chew | Chew | Eat |
| Sniff | Sniffing | Micro_move |
| RemainRU | Rearup | Rear |
| RemainPR | Rearup | Rear |
| RemainHV | Hang | Hang |
| RemainHC | Hang | Hang |
| RemainLw | RemainLow | Micro_move |
| WalkSlow | Walk | Walk |
| No.Data |  |  |
| Drnk.S2. | Drink | Drink |
| Drnk.S3. | Drink | Drink |
| Eat.Z2. | Eat | Eat |
| Eat.Z3. | Eat | Eat |

#### Analysis details

The software reads the **minute summary** file created by the HCS software or creates a new one from the raw behavior sequence data or the hourly summary data. In the latter case, the hour value divided by 60 is used for each minute of that hour. The software adds a column that indicates the time to the light-off event (“bintodark”) and what the light condition was (DAY or NIGHT). This is calculated from the start of the experiment in the experiment metadata (which could be read from the name of the video file coming from the HCS software package) and the light/dark cycle information obtained from the lab metadata file.

We used the information delivered by CleverSys to derive **categories** from the raw sequence files code, and obtained 38 categories (the distance traveled on the x axis was not considered a behavior category, ; No.Data and Arousal were discarded; and six different drink and eat categories were pooled into two). The software pools these 38 categories into 18 (Berlin categories) or 10 (Jhuang categories) using the “grouping variables.r” code (Table 2). The data records were typically from experiments lasting slightly less than 24 h. Nine different **time windows** were defined, with the last three windows overlapping with the first six; see Figure 2.

**Figure 2.**
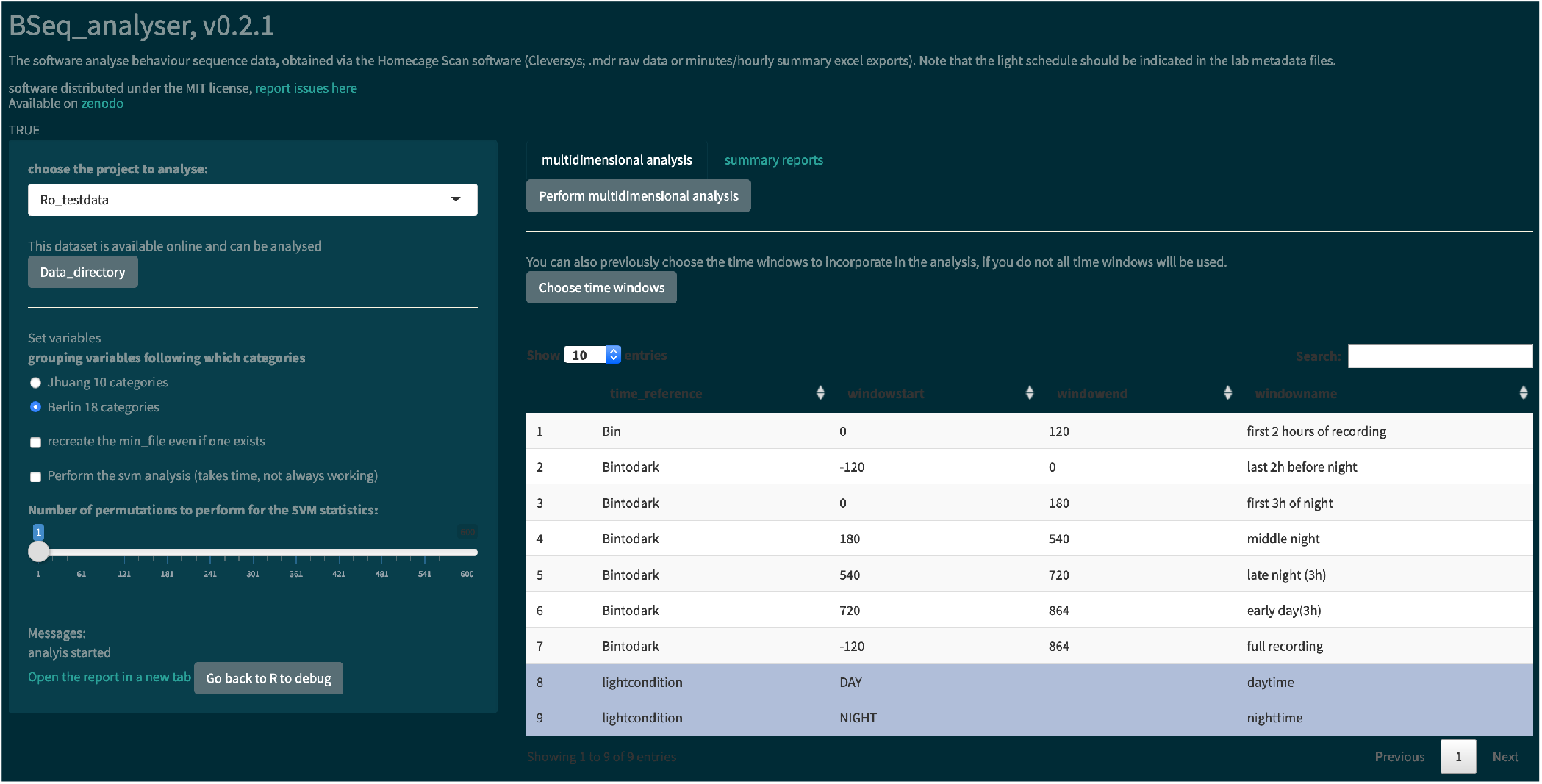
Screenshot of the Bseq analyser Shiny app for data analysis. In the left panel, the user indicates the data and variables to use. He or she can also indicate where to find the data that are not published online (by clicking the “Data directory” button). The left panel also presents some messages and a link to the report. The main panel has two tabs, one for performing the multidimensional analysis (with a prior choice of time windows or not) and one for creating hourly summary plots of each behavior category.

Then, the square root of the proportion of time spent performing each behavior during each time window is calculated. This **data transformation** makes the data more normally distributed and allows for a better analysis in a multidimensional space, but it does not require non-null values like log transformation. This derived data set is the multivariate dataset (called “Multi datainput”), which can contain up to 162 variables per subject (18 behavior categories times nine time windows).

#### Multivariate data visualization and analysis

The software uses a double RF analysis to select 8–20 variables, which are used as input for ICA. The first RF selects the best 20 variables, whereas the second RF is performed using only these 20 variables. The best eight variables or all variables with a Gini score above 0.95 are kept for the ICA and listed in the report. The data are then plotted according to the first three components of the ICA, resulting in a three-dimensional plot.

Then, the software performs a statistical analysis using a **PCA** on the multivariate data set, and then plots the first component and performs a statistical analysis of this first component over groups. Finally, the user can choose (via the “Perform the multidimensional analysis (takes time)” button) to perform a **machine-learning analysis** based on a support vector machine approach using a radial kernel. We also attempted an L1-regularized regression, modifying the code used in Steele et al. [2007], obtained from Prof. King. The models were used to predict the experimental group of the data not used for training. The software used two different **validation techniques**. For data sets with fewer than 15 animals per group, a two-out validation strategy is used, whereas the software uses a completely independent test data set when the sample size exceeds 15 (see the analysis details.md text delivered with the software for details). The software reports the kappa score as a measure of model accuracy. For the statistical analysis, the same machine-learning code is run on the same data but after a randomization of the group (permutation). This provides us with a cloud of accuracy results that can be used to perform a binomial test, which in turn provides us with a p-value that indicates whether the model can predict the experimental group at a level above chance.

## RESULTS

### Data integration

In order to facilitate the analysis of data from different sources, we proposed a format for organizing the data (behavior sequence or binned summary data) and the metadata (information about the experiment, the lab, and the animals), such that the R Shiny apps can access the different files automatically. Critically, this format does not require any file to be renamed, but it does include file names in the experiment metadata. An extra main and public metadata file reports information about the project, its contributors, and the placement of the other files (see Fig. 1), making the data FAIR [Group, 2014].

We have also provided a walk-through (available at metadata/information/Readme.md) and a Shiny app to facilitate new data integration. The app tests and uploads the project metadata to the master metadata file online (which can then be read by the second app devoted to data analysis). The process lasts for approximately 1 hour once all information have been gathered. The data files themselves did not require any modification. We obtained data of different quality and formats from different labs. The software deals only with files output from the HCS software package (CleverSys Inc.) thus far (the raw behavior sequence [.mbr file] or the minute or hour binned data summaries).

We provided and used a data set produced in Berlin of 11 wild-type female mice recorded twice (at the age of 3 and 7 months, respectively) for approximately 24 hours, and published data obtained from Andrew Steele [Steele et al., 2007, Luby et al., 2012b]. Other data sets were tested but the data were not made public. The sample size was decided independently of this study and one animal was excluded because the data for one time point were not available. Mice were tested in the same order at the two time points, and were subjected to other behavioral tests in the 4-month time period between the two home cage monitoring events.

### Data analysis

We used data obtained using the HCS software with 11 wild-type mice recorded twice for approximately 24 h. The data were grouped following the age of the animal (young or old) at the time of the recording (available under Ro testdata project). One Excel export file was corrupted (animal 279, first test), whereas the data of one animal was inconsistent (animal 25, second test: the raw data and the exported data did not correspond). Animal 25 was removed from the analysis by modifying the metadata file, which contained an “exclude” column.

The second R Shiny app was used to analyze the data, as shown in Figure 2. In the left panel, the user must choose variables: the project to analyze, the behavior categorization to use (Table 2), whether to recreate the minute summary file from the raw data, whether a machine learning analysis should be performed, and the number of permutations to perform (if machine learning analysis is performed). The users might then choose which time windows to incorporate in the analysis. They then press the “Perform multidimensional analysis” button and wait until the html report is produced and presented on screen. We performed thise analysis once with the corrupted data from the excel summary files (Fig. 3) and once with a corrected export of the raw data (see whole report at https://doi.org/10.6084/m9.figshare.6724547.v3).

**Figure 3.**
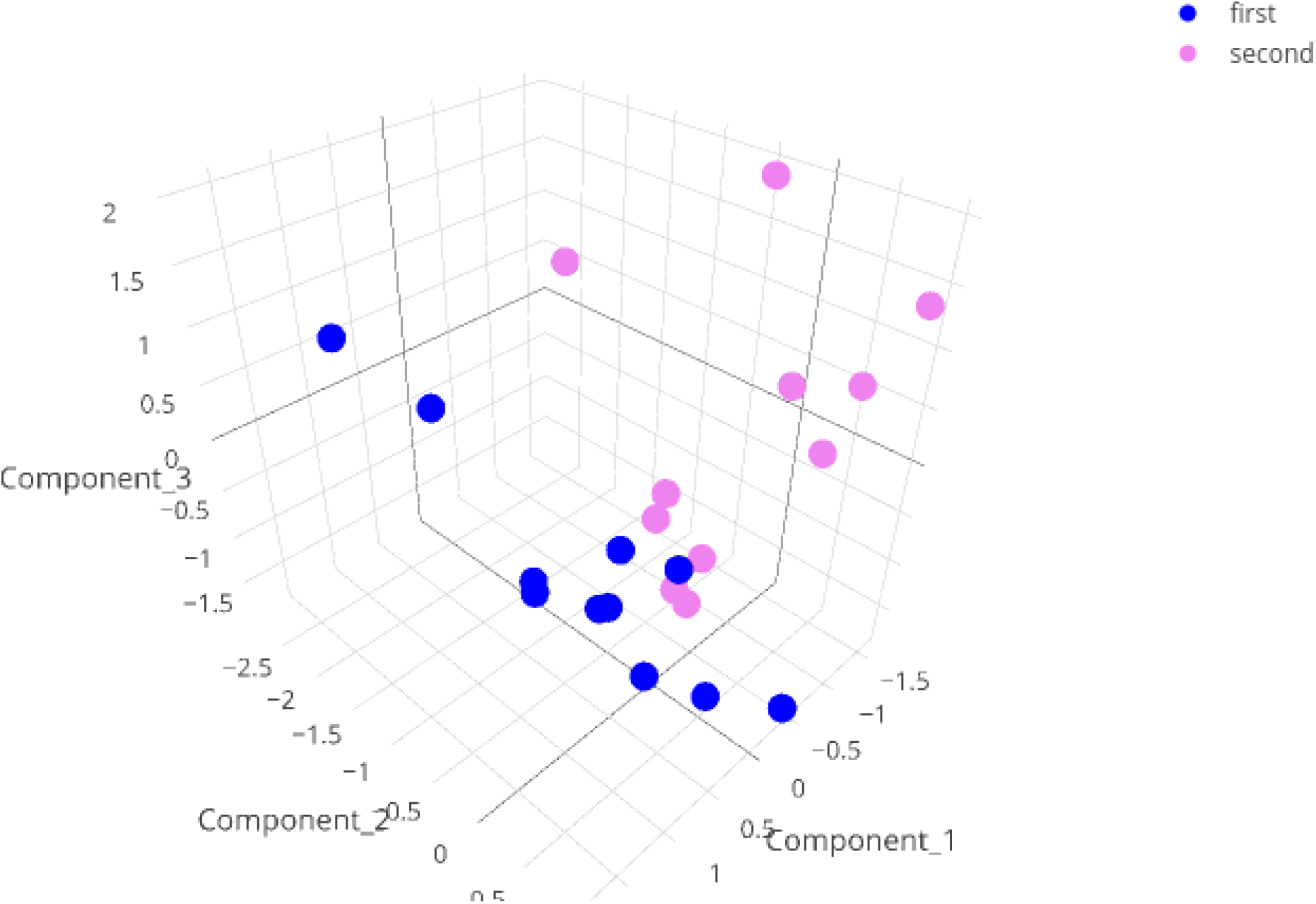
3D representation of the results of the ICA on the test data with a minute summary as the primary data. Note that the data with corrupted entries (animal 279, first test) does not show up as an outlier in this graph.

The software read and transformed the data according to the information given in the metadata and the variables selected. It reports a data analysis using a PCA, present the results of RF analysis and visualize the data in 3 dimensions. (for details, see the Materials and Methods section and the code itself). The html report is saved on the hard disk (see Fig. 2) and can be directly opened in the browser app. Noteworthily, the PCA was effective at separating the two experimental groups (nonparametric statistical test on the first component [p = 0.00067; the effect size was large: Z/square(n) = 0.76]).

For the data visualization, a random forest algorithm was used to choose the 8–20 variables that were the most effective at separating the different animal groups. An independent component analysis (ICA) was then run on these variables and the data were plotted in two or three dimensions. When we performed this analysis including the corrupted file, the data point was surprisingly not an extreme value (see Fig. 3, interactive at https://plot.ly/~j_colomb/39/).

In the second tab of the app, hourly summaries of the percentage of time spent performing each behavior are provided (time is synchronized to the light-off event). Although it can be directly seen in the app (plot by plot), a pdf file with all plots is also produced. Figure 4 presents an example of a 24-h summary plot obtained using the “walking” behavior category.

**Figure 4.**
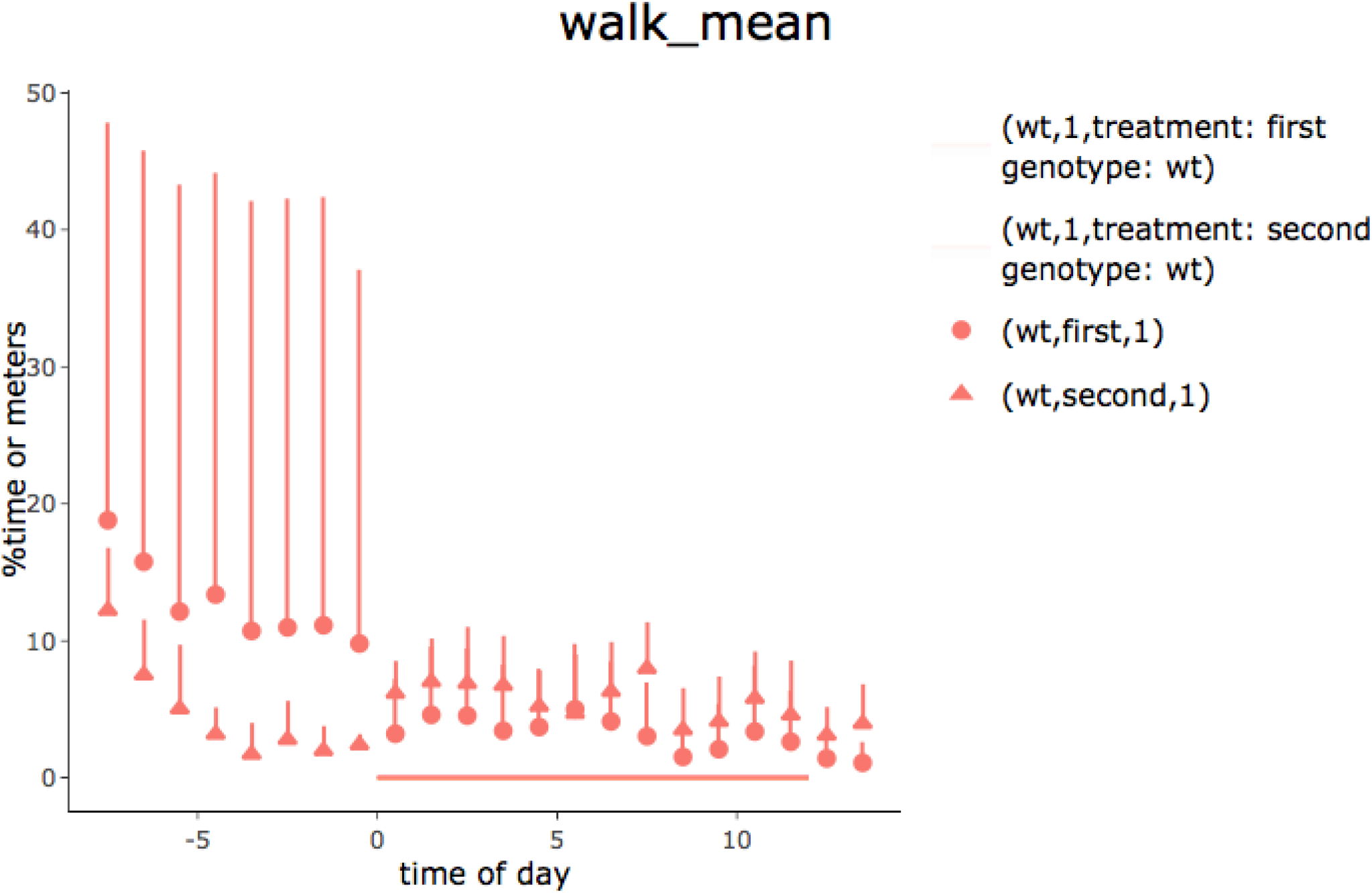
Time spent walking for young (first) and old (second) animals observed for 24 h (lights went off at 0 h). Recordings began immediately after the animals had been placed in the observational cages. Data are means and standard deviation expressed as percentages of time spent walking. The increase in activity at the start of the test appeared to wear off faster in old animals. In order to work for different types of grouping, the legend shows genotype and treatment conditions and is not optimized.

**Figure 5.**
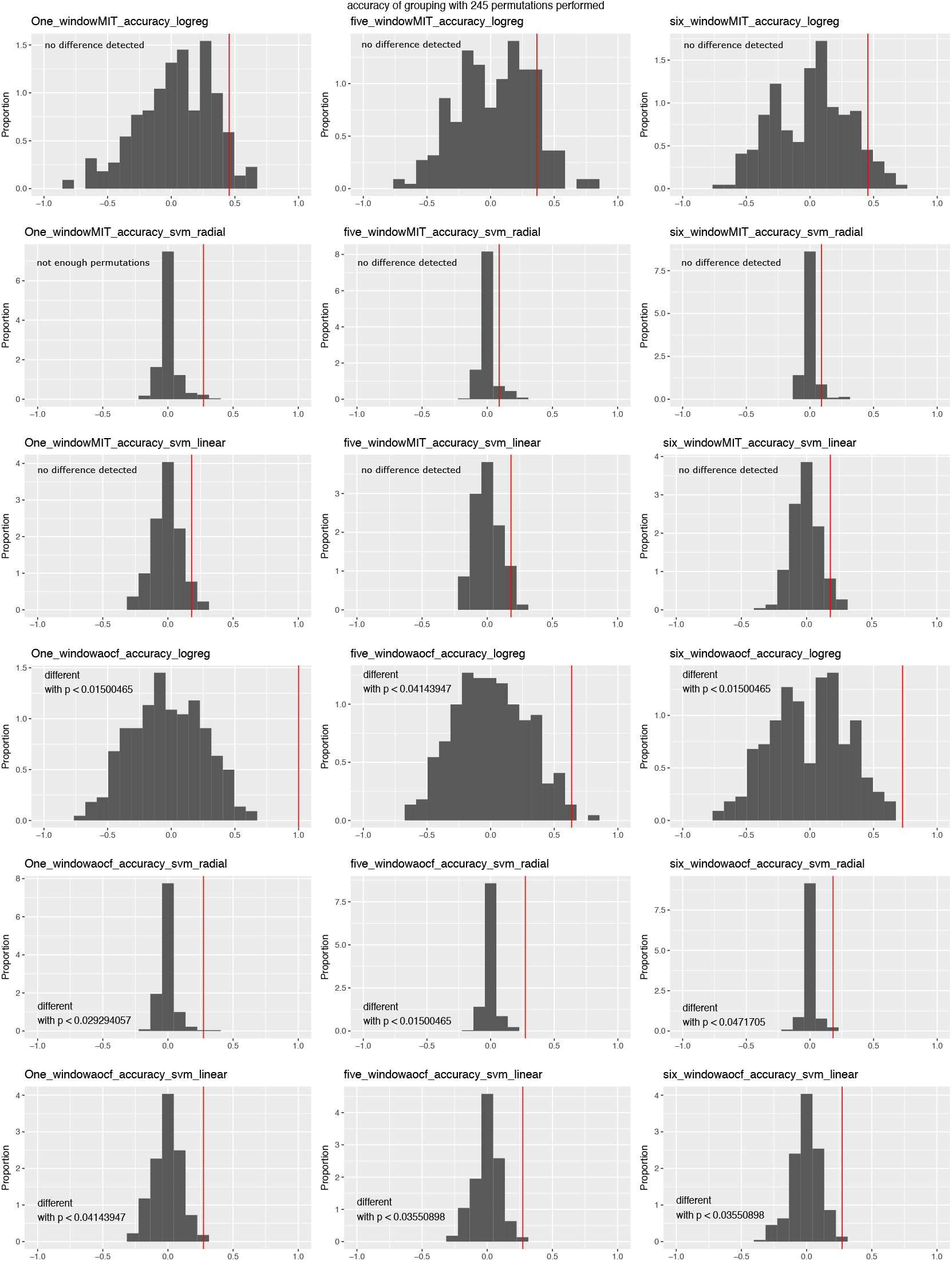
Accuracy of machine-learning algorithms in predicting data group membership in the test data, using two-out validation. The red line represents the accuracy when the real groups were used, whereas the distribution represents the accuracy obtained when data group membership was randomized prior to the analysis. Graphs are grouped according to the number of time windows used in the columns (one, five, or all six windows; see text), categorization of the behavior (first three rows: Jhuang categorization, last three rows: Berlin categorization) and the machine-learning algorithm (L1-regularized regression: rows 1 and 4; SVM with radial kernel: rows 2 and 5; SVM with linear kernel: rows 3 and 6); p-values were obtained through confidence intervals for binomial probability analysis.

### Machine learning analysis

The software predicts group separation using a multidimensional analysis. In addition to PCA, it might then use a supervised machine learning (support vector machine [SVM] using a radial kernel) approach to separate the two groups. Noteworthily, the software can also deal with three different experimental groups, but no more (the data had to be split with pairs of groups and an analysis was performed for each pair). The SVM is trained on part of the data. The model is then used to predict the group membership of data not used to train the model. The kappa score gives an indication of the effectiveness of the model, which itself indicates how easy the two groups of data can be separated. The software uses either a two-out validation strategy (as in Steele et al. [2007])if there are fewer than 15 animals per group, or an independent test data set otherwise. The whole process is repeated after permuting the group membership of the train data set. A binomial test compares the actual accuracy with a cloud of accuracy values obtained after many permutations, thus calculating a range for the p-value. The number of permutations is reported with this estimated p-value.

In order to test the efficacy of the approach, we ran the analysis using different variables with our test data set. This was performed with version v0.1.1-alpha of the software and with the two corrupted files for animals 25 and 279. While the PCA could tell the experimental groups apart (data not shown), the machine-learning approach was not as effective. We performed analyses over three time window variations: one time window (from 2 h before lights off to the end of the recording), five time windows (first 2 h of recording, last 2 h before nightfall, first 3 h of the night, last 3 h of the night, first 3 h of the second day), or six time windows (all of the windows described above). We ran the analysis using both behavior categorizations. Since the number of animals was low (11 per group), we used the two-out validation procedure. The algorithm could tell the two groups apart when the Berlin categorization was used, but not when the Jhuang categorization was used, irrespective of the time window combination or algorithm used (Fig.5). When the same analysis was performed with corrected data, the latest code and the one time window in the Berlin categorization seemed to provide an even worse success rate for the SVM approach https://doi.org/10.6084/m9.figshare.6724547.v3.

### Working with hourly summaries and raw data

The software could also use hourly summaries as primary input data (they are the only data available in the Steeleo7 HD data set). In this case, a minute summary was produced by dividing the hourly value by 60. The synchronization with lights off between experiments was not precise in those cases, but a rough analysis of the output revealed that this had only a minor effects on the whole analysis (https://doi.org/10.6084/m9.figshare.6724604.v1).

In general, we recommend exporting minute summaries from the HCS software for new experiments (to obtain the distance traveled) but using the raw data for analysis. Indeed, the distance traveled per minute cannot be calculated with our software. However, the created minute summary file is more robust than that in the HCS software; specifically, some behavior events were sometimes not taken into account, and in one case the HCS export function failed completely.

Remarkably, using the raw data as the input allows for a more complex analysis of the data. One can, for instance, analyze the transition between different behaviors. For example, we showed which behavior was performed before and after “land vertical” events, merging our two experimental groups (Fig. 6). While landing occurred after hanging behaviors as expected, the animals started to either rear again or engage in sniffing or eating behaviors, but rarely started to walk directly after a landing. In addition, the “hanging vertical from the rear up” behavior notably did not follow a rear up behavior in these cases.

**Figure 6.**
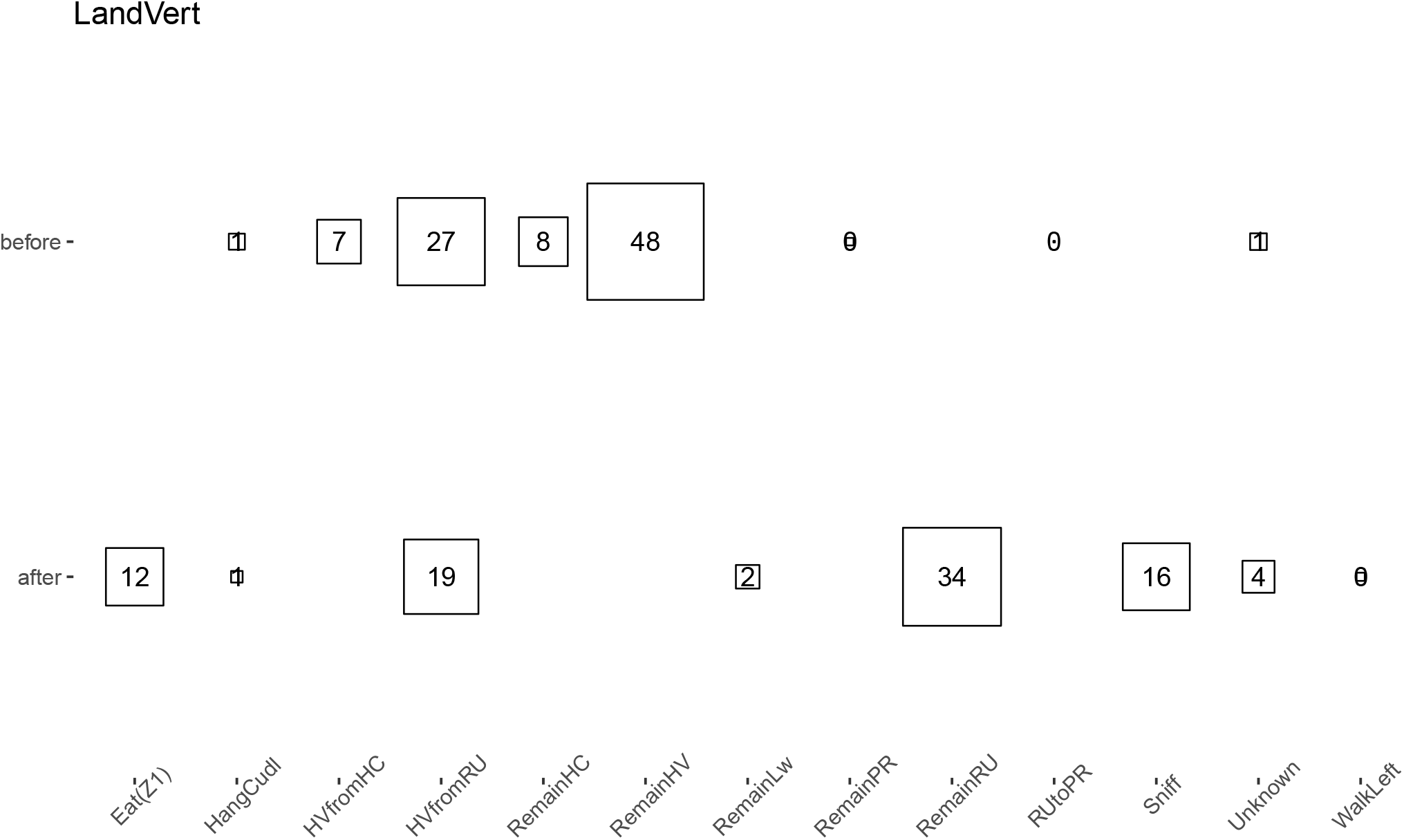
Average percentage of time that a behavior (the original HCS behavior categories were used) appeared just before or after a “land vertical” behavior. The eight behaviors with the highest median proportion are shown; squares and numbers represent the mean percentage. Similar numbers were obtained when taking the median.

### Meta-analysis

In order to test the re-usability of our data and code, we performed a meta-analysis using data from different projects. We read all data at our disposal for wild-type animals. We then performed the usual analysis with all nine time windows, followed by a PCA (we could not include the Steele07 HD data because the birth date of the animals was not provided, and also because the seventh time window did not have data). We plotted the first PCA component against the age of the animal, adding the genotype as an additional variable (Fig. 7). The results suggested that both age and genotype might affect mouse behavior in the home cage.

**Figure 7.**
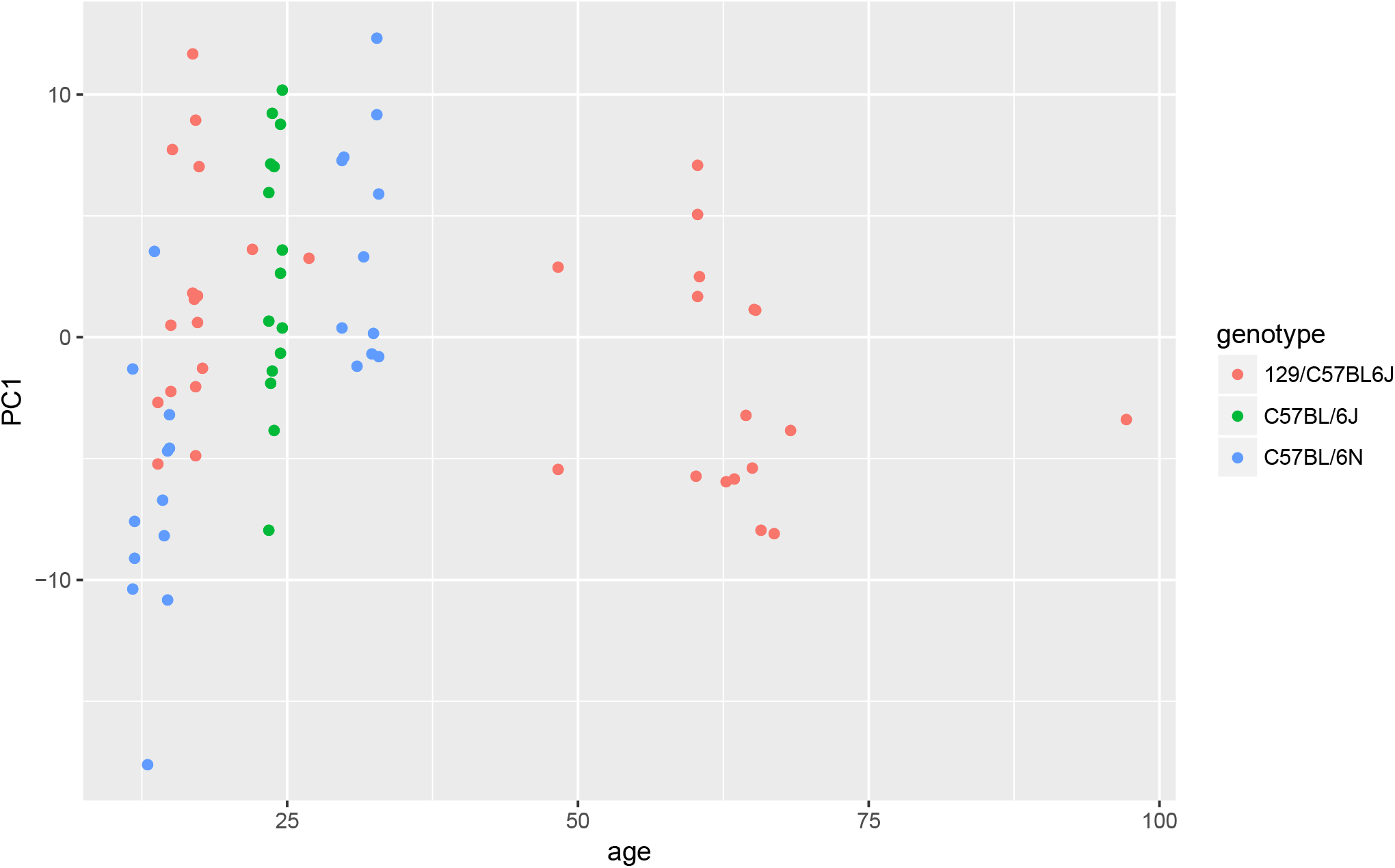
Effect of age and genotype on behavior phenotype as summarized by the first component of a PCA. The data suggested that both genotype and age could affect animal behavior.

## DISCUSSION

### FAIR data per default

Using relatively simple tools (R and spreadsheets) and common platforms (GitHub, OSF, and Zenodo), we combined data analysis and data “FAIRification” into one workflow. On top of metadata necessary for the data analysis, we ask the users to provide general information about the experiment, and strongly encourage them to publish this particular piece of metadata through one of our apps (Fig. 1). This creates an open repository for home cage monitoring metadata in a spreadsheet form (https://osf.io/myxcv/). Users may choose to keep the data private, but even unpublished data is in a state to be shared easily.

Home cage monitoring experiments lead to videos that are analyzed to produce a time stamped sequence of recognized behaviors. By combining these data with metadata (which provide information about the experiments and the experimenters in a computer-readable form), one can produce interesting visualizations and analyses, especially if the raw data (in this case an .mbr text file) are provided. We encourage users to avoid using the Excel summary files produced by the proprietary software, but rather to start the analysis from the raw data. Doing so will make the analysis more robust: data from different software may be included more easily, and one avoids problems created in closed access export functions. In particular, we encourage users not to include the distance traveled variable in the analysis, as its spread differs from the other variables (percentage of time spent performing a behavior) and thus including it in a multidimensional analysis may cause problems.

In order to best test the data and code re-usability, we performed a meta-analysis (Fig. 7). We pooled all data available to us, filtered those from wild-type strains, and asked whether animal age or genotype had the most influence on the animal behavior. While the amount of data available to us proved insufficient to answer the question, the analysis could be performed with few issues.

### Data visualization and analysis

The software aligns the data to the light/dark cycle, and then cleans the data to only keep data points where all subjects have provided data, thus ensuring that each sample is of equivalent valence for the analysis. Such data cleansing has been absent from most analyses published to date, although it might be crucial for spotting specific effects at the time of the light/dark switch. We also implemented different time windows to create specific variables along the day/night cycle, in order to detect differences that could be overseen with a 24-h summary analysis. We are confident that the software represents progress toward a cleaner and more detailed analysis of behavior sequence data.

In addition, the use of the software would prevent harking [Kerr, 1998]. To illustrate this, we shuffled the grouping of the test data before performing the analysis (data not shown). We observed that the 3D data visualization still showed some differences between the groups. This was expected because the RF analysis is meant to look for the cause of differences and would find some in a data set with 162 variables. The summary analysis (which corresponds to the type of analysis usually performed) can still reveal some apparent differences for some behavioral traits – differences that could be claimed to be statistically significant if one does not correct for multiple testing. However, the PCA clearly indicated that a difference between the two artificial groups of data could not be detected statistically, as expected.

We made quite an effort to implement and test a machine-learning approach, with the idea being that a PCA may miss existing differences due to high variability for some variables inside groups. However, our analysis revealed that this approach seems to be ineffective with our type of data (Fig. 5). In particular, the analysis revealed that the distribution of the accuracy of predictions in randomly permuted groups varied greatly between algorithms, which questions the approach used by Steele et al. [2007].

### An open source proof of concept

By using a GitHub workflow and an open-source programming language (R), providing Shiny apps for use by non-coders, and implementing metadata in simple spreadsheets that are easy to read and write, we hope to reach the growing community of researchers who are dealing with behavioral sequence data. The software is intended for non-computer-scientist researchers to read and extend, and therefore, it has been kept simple. While we have provided extensive comments, including dependencies, as well as a hierarchy of code files to facilitate code reading, we did not use functions nor implement tests. Similarly, the experiment metadata are provided in spreadsheets, a practical solution that could be implemented without much effort. We believe that the implementation of more complex data format would be counterproductive at his stage.

The analysis runs identically on the Shiny app or when variables are provided in a code file, so debugging and extension creation can be performed without the need to care about the difficulties of Shiny apps debugging. We used that approach to perform a quick analysis of the behavior transition in our data set. Our results demonstrated the potential of this approach both for spotting limits in the video analysis software (e.g., inconsistent sequences) and for creating new, more detailed analyses based on the behavior sequence itself.

### Mouse behavior

As expected, mouse behavior differed in the second session compared with the first session, which was detected by a PCA. An explorative look at the data suggested that mice are more active at the beginning of their first session, during the day, confirming that the use of different time windows is beneficial for the analysis of the data. Our meta-analysis also suggested that both age and genotype influence mouse behavior.

## CONCLUSION

We have presented several open-source Shiny apps that allow the archiving, visualization, and analysis of long-term home cage video monitoring experiments. This report is a proof of concept for workflows allowing both data analysis and publication. The analysis tool by itself should be helpful for the analysis of behavioral sequence data. It cleanses the data before analysis and provides an easy way to test for group effects while avoiding harking and p-hacking. We hope that the community will increase the amount of data openly available as well as expand the software in novel ways for analyzing behavioral sequence data.

## APPENDIX

## Competing interests

none declared

## Grant information

This work was funded by the Deutsche Forschungsgemeinschaft (DFG, German Research Foundation), SFB 1315, Project-ID 327654276, and EXC 2049: NeuroCure, Project-ID 390688087. We acknowledge support from the Open Access Publication Fund of Charité – Universitätsmedizin Berlin.

## Acknowledgements

We wish to thank the members of the lab and the AOCF team: Andrei Istudor for his suggestion to integrate all used variables in the software report; Patrick Beye for the discussion on the design of the analysis; Melissa Long for performing the home cage monitoring experiment and for helping with the creation of the Metadata; and Vladislav Nachev for the scientific input and discussion. In addition, we wish to thank Prof. Steele and Prof. King for their fruitful discussions and access to their code and data, and also CleverSys Inc. for their help with decoding the HCS outputs.

## Notes

https://github.com/jcolomb/HCS_analysis

### Competing Interest Statement

The authors have declared no competing interest.

